# Cortico-hippocampal interactions underlie schema-supported memory encoding in older adults

**DOI:** 10.1101/2024.09.18.613755

**Authors:** Shenyang Huang, Paul C. Bogdan, Cortney M. Howard, Kirsten Gillette, Lifu Deng, Erin Welch, Margaret L. McAllister, Kelly S. Giovanello, Simon W. Davis, Roberto Cabeza

## Abstract

Although episodic memory is typically impaired in older adults (OAs) compared to young adults (YAs), this deficit is attenuated when OAs can leverage their rich semantic knowledge, such as their knowledge of schemas. Memory is better for items consistent with pre-existing schemas and this effect is larger in OAs. Neuroimaging studies have associated schema use with the ventromedial prefrontal cortex (vmPFC) and hippocampus (HPC), but most of this research has been limited to YAs. This fMRI study investigated the neural mechanisms underlying how schemas boost episodic memory in OAs. Participants encoded scene-object pairs with varying congruency, and memory for the objects was tested the following day. Congruency with schemas enhanced object memory for YAs and, more substantially, for OAs. FMRI analyses examined how cortical modulation of HPC predicted subsequent memory. Congruency-related vmPFC modulation of left HPC enhanced subsequent memory in both age groups, while congruency-related modulation from angular gyrus (AG) boosted subsequent memory only in OAs. Individual differences in cortico-hippocampal modulations indicated that OAs preferentially used their semantic knowledge to facilitate encoding via an AG-HPC interaction, suggesting a compensatory mechanism. Collectively, our findings illustrate age-related differences in how schemas influence episodic memory encoding via distinct routes of cortico-hippocampal interactions.

## Introduction

Compared to young adults (YAs), older adults (OAs) are impaired in memory for novel events (*episodic memory*). In contrast, memory for general knowledge of the world (*semantic memory*) is preserved or even enhanced in OAs (Levine et al. 2002; Park et al. 2002). A core component of semantic memory is the knowledge of schemas, which are extensive relationships among concepts, formed by abstracting commonalities and associations across multiple experiences; in turn, schemas provide scaffolds unto which novel information can be assimilated, enhancing subsequent cognitive processing and memory (Ghosh and Gilboa 2014; Umanath and Marsh 2014; Varga et al. 2022). There is abundant behavioral evidence that OAs can take advantage of their preserved schematic knowledge to ameliorate their episodic memory deficits (Umanath and Marsh 2014). The current fMRI study investigates the neural mechanisms of this phenomenon.

The finding that schemas facilitate the acquisition of novel episodic information has been replicated many times using a variety of different paradigms and stimuli, including pre-existing knowledge of grocery item prices (Castel 2005; Whatley and Castel 2022) and home interior layouts (Hess and Slaughter 1990; Arbuckle et al. 1994), as well as rules and associations acquired during the experiment (Wagner et al. 2015; Brod et al. 2017; De Brigard et al. 2017; Greve et al. 2019). In a popular paradigm, participants view pictures of scenes paired with objects, which are either (1) expected in the scene (*congruent*; e.g., garden-flower), (2) neither expected nor unexpected (*neutral*; e.g., garden-statue), or (3) unexpected in the scene (*incongruent*; e.g., garden-television) (van Kesteren et al. 2013; van der Linden et al. 2017; Greve et al. 2019; Quent et al. 2022). Memory is typically better for congruent than neutral objects (*congruency effect*) and is sometimes also better for incongruent than neutral objects (*incongruency effect*) (Chen et al. 2022; Quent et al. 2022; Klever et al. 2023). Whereas the congruency effect is assumed to reflect a semantic elaboration (the congruent object is successfully encoded because it fits with the schema), the incongruency effect is assumed to be attentional, owning to novelty and prediction error (Greve et al. 2019). Consistent with the latter, the incongruency effect is more common when the incongruent object is presented within the scene (e.g., a chainsaw on top of a kitchen counter), thereby generating greater surprise (Friedman 1979; Goodman 1980; Wynn et al. 2020). In the current study, the object is presented after the scene, and hence, we focus primarily on the congruency effect and its neural mechanism.

As noted above, episodic memory deficits in OAs are attenuated when the learning environment encourages the use of pre-existing schemas (Park and Reuter-Lorenz 2008; Zimerman et al. 2011). The possible explanation of this presumptive “schematic support effect” is that OAs take advantage not only of their relatively preserved semantic knowledge (Haitas et al. 2023) but more critically of the rich schematic knowledge they accumulated and consolidated during a lifetime to facilitate successful encoding and retrieval of new information within appropriate models of the world (Craik and Bosman 1992; Chen et al. 2022). Consistent with this idea, the congruency effect in the scene-object paradigm (Brod and Shing 2019; Chen et al. 2022) and other paradigms (Castel 2005; Badham et al. 2016; Mohanty et al. 2016; Whatley and Castel 2022) tends to be larger for OAs than YAs. However, while the function of regions underlying schema processing has been a focus of research for several years (reviewed below), the neural mechanisms of the age-related schematic support effect are largely unknown, and the focus of the current study.

The evidence on the neural mechanisms of schema processing and their impact on episodic encoding comes primarily from fMRI studies with YAs and primarily implicates four brain regions: hippocampus (HPC), ventromedial prefrontal cortex (vmPFC), anterior temporal lobe (ATL), and angular gyrus (AG). HPC is critical for the encoding of new items and the formation of novel associations (Teyler and DiScenna 1986; Squire 1992; Treves and Rolls 1994; Davachi and Wagner 2002; Ranganath et al. 2004), both of which are essential functions for schema-related processes. The hippocampal memory indexing theory (Teyler and DiScenna 1986) suggests while heterogeneous informational contents are stored in disparate cortical regions, HPC caches the links to those representations and reactivates them during subsequent recollections. Corroborating this idea, previous research has found that hippocampal interactions with multiple cortical regions, including the primary visual cortex, inferior frontal gyrus, and AG, supported subsequent memory (Huang, Howard, et al. 2024).

VmPFC has been strongly associated with processing schemas and their impact on episodic memory (van Kesteren et al. 2012; Gilboa and Marlatte 2017). FMRI evidence indicates that vmPFC instantiates and maintains appropriate schemas during various tasks (van Kesteren et al. 2010; van Kesteren et al. 2013; Gilboa and Moscovitch 2017; Masís-Obando et al. 2022). Moreover, vmPFC’s functional coupling with HPC has been suggested to underlie the integration of isolated memory representations and the formation of new schemas (van Kesteren et al. 2010; Zeithamova et al. 2012). However, the literature is mixed about whether vmPFC-HPC connectivity is associated with the congruency effect, the incongruency effect, or both. Whereas some evidence links vmPFC-HPC connectivity to better memory (across-participants) for congruent information (Liu et al. 2017), other findings link it to better memory (within-participants) for incongruent but not congruent information (van Kesteren et al. 2010; Bein et al. 2014). Thus, further research on vmPFC-HPC connectivity during schema-related learning is warranted.

ATL is assumed to be an amodal hub of conceptual knowledge and semantic memory, an idea supported by patient, fMRI, and modeling evidence (Patterson et al. 2007; Lambon Ralph et al. 2017). ATL is also an important node in the anterior temporal network and connected with anterior HPC for processing semantic information (Ranganath and Ritchey 2012; Ritchey et al. 2015). Although ATL has not been the focus of typical schema paradigms, such as scene-object paradigm, it has been associated with a form of schema-based learning known as fast-mapping (Atir-Sharon et al. 2015; Merhav et al. 2015; Farahibozorg et al. 2021; Zaiser et al. 2022).

Finally, AG is a core semantic region and has been also associated with schema processing and its impact on new learning. AG acts as a convergence zone that binds together multimodal percepts (Bonner et al. 2013; Bonnici et al. 2016; Humphreys et al. 2021), as well as integrates abstract concepts (Price et al. 2015; Kuhnke et al. 2023). Indeed, the neural representations of distinct components of a schema were found to converge in the AG, both after consolidation and during the transfer to new related tasks (Wagner et al. 2015). An alternative idea is that AG does not store semantic representations but rather it controls access to semantic representations stored in other regions, such as ATL (Lambon Ralph et al. 2017). AG has extensive structural and functional connections with HPC (Rushworth et al. 2006; Uddin et al. 2010; Tambini et al. 2018; Hermiller et al. 2019), with AG-HPC functional connectivity parametrically increasing with stronger schema congruency (van der Linden et al. 2017). The AG-HPC interaction was also found to be mnemonically relevant, such that the coordinated representation of semantic information in these two regions at encoding predicted subsequent memory of object concepts (Huang, Howard, et al. 2024).

Thus, despite abundant behavioral evidence of the schematic support phenomena in OAs, our neuroscientific understanding of schemas and their impact on memory have been limited to YAs. To our knowledge, only one study (Brod and Shing 2019) has directly investigated the neural correlates of schematic support in OAs.

Focusing on vmPFC activity during encoding, the authors found a cluster that showed a greater subsequent memory effect (i.e., “Remembered > Forgotten”) for congruent than incongruent trials, although no age effect was found. Moreover, the study did not investigate other relevant semantic regions, such as AG or ATL, and was limited to univariate activity results. In contrast, in the current study, we investigate how these different cortical regions modulate episodic encoding processes in HPC.

The current fMRI study consists of three sessions. In Session 1, younger and older participants were shown a series of everyday objects once, for familiarization. In Session 2, a week later, they explicitly associated each object with a real-world scene by rating the likelihood of finding the object there, providing a subjective rating of schema congruency. Finally, in Session 3, a day later, participants’ memory for the objects was tested with an old/new recognition task (see **Figure 1**). At the behavioral level, we predicted that both YAs and OAs would show a congruency effect, and that the congruency effect would be larger for OAs than YAs. To investigate the neural mechanism of this effect, we use a modified psychophysiological interaction (PPI) analysis to explicitly examine how much vmPFC, AG, and ATL modulate HPC encoding processes, enhancing subsequent memory for congruent trials. We predicted that in these regions, the schema-related modulation of HPC encoding processes would be stronger for OAs than YAs. Such result would explain the predicted behavior and would provide a potential neural mechanism whereby OAs compensate for episodic encoding deficits by over-relying on semantic schemas.

**Figure 1.**
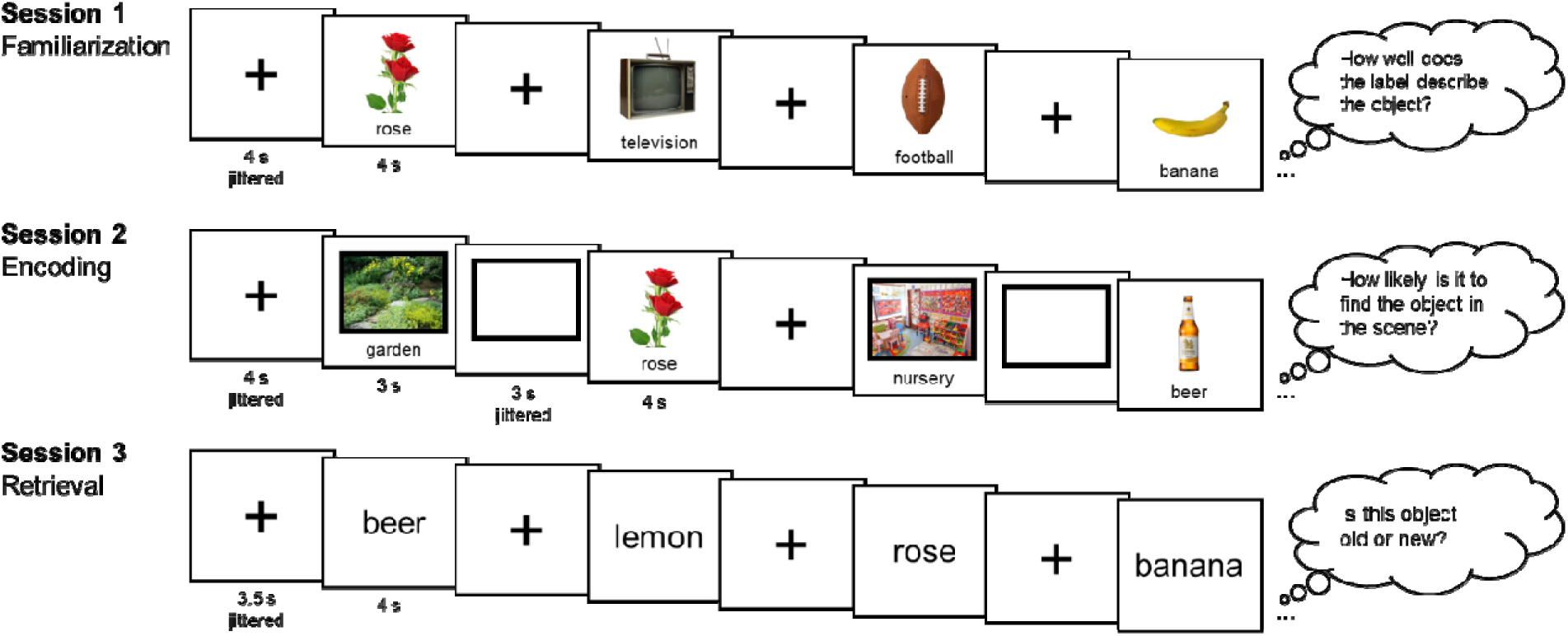
Experimental design. The experiment consisted of three sessions. In Session 1 (Familiarization), images of unique everyday objects were presented serially, and participants rated how well the given label described the object from 1=“does not describe the object” to 4=“exact description”. In Session 2 (Encoding), a week later, the objects from Session 1 were presented again, each preceded by a real-world scene. For each object, participants rated how likely it is to find the object in the scene from 1=“very unlikely” to 4=“very likely”. For each participant, encoding trials we divided into three sets with different distinct levels of schema congruency based on the subject ratings. Finally, in Session 3 (Retrieval), a day later, participants completed a conceptual memory task, where they were presented with labels of previously seen objects intermixed with the labels of unrelated new objects, and they rated the oldness of the objects from 1=“definitely new” to 4=“definitely old”. Participants also completed a perceptual memory task, which was not analyzed in this study.

## Materials and Methods

### Participants

A total of 38 YAs and 38 OAs participated in this study voluntarily for monetary compensation. To be eligible, participants must be between 18 and 30 years of age (YA) or between 65 and 85 years of age (OA), must be a native or fluent speaker of English, must have no history of significant neurological or psychiatric conditions, and must not be taking medications that affect cognitive function or cerebral blood flow (except antihypertensive agents). OAs were further screened for cognitive impairment with the Montreal Cognitive Assessment (MoCA) (Nasreddine et al. 2005) and must obtain a minimum score of 23 (Rossetti et al. 2011), which roughly equals a Mini-Mental State Examination (MMSE) score of 28 (Trzepacz et al. 2015). All participants provided written informed consent prior to participation. Four YAs and eight OAs did not complete the study and were excluded from analysis. Two YAs and two OAs were further excluded due to very low object recognition memory (see *Analysis plan – Quality checks*). The final sample included 32 YAs (age range = [19, 29], M = 23.12, SD = 3.31; 20 females, 12 males) and 28 OAs (age range = [65, 82], M = 71.79, SD = 4.43; 19 females, 9 males).

### Experimental design

The study protocol was approved by the Duke University Health System Institutional Review Board, and all methods were performed in accordance with the relevant guidelines and regulations. The experiment consisted of one session of cognitive assessment and three fMRI sessions (see **Figure 1**). Immediately before the start of each scanning session, participants were instructed on the corresponding task and completed a short practice run. Session 1 consisted of three scanning runs of 38 trials, and each run lasted approximately 5 min 24 s. Participants viewed images of 114 unique everyday objects on a white background and rated how well the given label describes the object on a 4-pt scale (1=“does not describe the object”, 2=“poor description”, 3=“good description”, 4=“exact description”). The main purposes of this task during familiarization were to ensure participant engagement with the meaning of the stimuli, and to ensure the quality of the verbal labels (mean = 3.60). Object presentations lasted for 4 s each and were separated by a jittered fixation cross that lasted for 4 s on average. After fMRI, participants completed the cognition battery from the National Institutes of Health (NIH) Toolbox (Denboer et al. 2014) in an adjacent room. The present study used participants’ crystallized intelligence as an index of general knowledge and semantic memory.

Session 2 occurred at least 7 days after Session 1 and consisted of three scanning runs of 38 trials, with each run lasting approximately 9 min 12 s. Participants viewed images of scenes of 114 unique real-world scenes and the 114 objects they viewed in Session 1. In each trial, participants were presented with a scene image for 3 s, a jittered empty box for an average of 3 s to indicate the continuation of the trial, and finally an object image for 4 s. The serial presentation of the scene and object allowed us to separate the neural activity of schema instantiation from that of object encoding. During object presentation, participants rated “how likely it is to find the object in the scene” on a 4-pt scale (1=“very unlikely”, 2=“somewhat unlikely”, 3=“somewhat likely”, 4=”very likely”). The pairing of scene and object was created such that, one-third of the objects would be considered congruent with the schema provided by the preceding scene (e.g., garden ➔ rose), one-third of the objects would be incongruent (e.g., bakery football), and the final one-third would be neutral (e.g., woods ➔ railroad). Three counterbalanced sets of unique scene-object pairings were created such that each scene and each object would appear in all three congruency conditions across participants. Trials were separated by a jittered fixation cross with an average of 4 s.

In Session 3, which happened one day after Session 2, participants were tested on their memory of the objects. The conceptual memory task consisted of three scanning runs of 48 trials, with each run lasting approximately 6 min 20 s. Participants were presented with 144 labels of real-world objects, amongst which 114 were labels for objects they had previously seen and 30 were unrelated novel distractors. While the word was on screen (4 s), participants were asked to respond whether the word referred to an old object or a new object (1=“definitely new”, 2=“probably new”, 3=“probably old”, 4=“definitely old”). Trials were separated by a jittered fixation cross that lasted for 3.5 s on average. Next, the perceptual memory task consisted of three scanning runs of 42 trials, with each run lasting approximately 5 min 36 s. Participants were presented with 126 images of real-world objects, amongst which 96 were old images that they had seen before (32 per congruency level), 18 were a different exemplar image of an old object (e.g., a different *rose* exemplar), and 12 were images of unrelated novel objects. While the image was on screen (4 s), participants were asked to respond whether the image was exactly old, a new image that depicted an old object, or completely novel. Trials were separated by a jittered fixation cross that lasted for 3.5 s on average. All analyses in the present study pertained to fMRI data from Session 2, congruency ratings from Session 2, and conceptual memory responses from Session 3. Data for the remaining parts of the experiment will be reported elsewhere.

### MRI data acquisition

MRI data were acquired on a 3T GE MR750 Scanner with an 8-channel head coil located in the Brain Imaging and Analysis Center at Duke University. Each MRI session began with a localizer scan, during which 3-plane (straight axial/coronal/sagittal) localizer faster spin echo images were collected. A high-resolution T1-weighted (T1w) structural scan image (96 axial slices parallel to the AC-PC plane with voxel dimensions of 0.9×0.9×1.9 mm^3^) was collected, followed by blood-oxygenation-level-dependent (BOLD) functional scans using a whole brain gradient-echo echo planar imaging sequence (repetition time = 2000 ms, echo time = 30 ms, field of view = 192 mm, 36 oblique slices with voxel dimensions of 3×3×3 mm^3^). Task instructions and stimuli were presented by the PsychToolbox program (Kleiner et al. 2007) and projected onto a mirror at the back of the scanner bore. Responses were recorded using a four-button fiber-optic response box. Participants wore earplugs to reduce scanner noise and, when necessary, MRI-compatible lenses to correct vision. Foam padding was placed inside the head coil to minimize head movement. We also collected resting-state images and diffusion-weighted images as part of a broader project, but those data will be reported elsewhere.

### MRI data preprocessing

fMRIPrep 23.0.1 (Esteban et al. 2019; Esteban et al. 2023) was used to preprocess structural and functional MRI data. Textual descriptions of preprocessing details generated by fMRIPrep are summarized below. T1w structural images collected across all MRI sessions for the same participant were corrected for intensity non-uniformity with ‘N4BiasFieldCorrection’ (Tustison et al. 2010) from ANTs 2.3.3 (Avants et al. 2008). The T1w-reference was skull-stripped with a Nipype implementation of the ‘antsBrainExtraction.sh’ workflow from ANTs, using OASIS30ANTs as target template. Brain tissue segmentation of cerebrospinal fluid (CSF), white-matter (WM), and gray-matter (GM) was performed on the brain-extracted T1w using ‘fast’ from FSL (Zhang et al. 2001). An anatomical T1w-reference map was computed after registration of T1w images using ‘mri_robust_templatè from FreeSurfer 7.3.2 (Reuter et al. 2010). Brain surfaces were reconstructed using ‘recon-all’ from FreeSurfer 7.3.2 (Dale et al. 1999), and the brain mask estimated previously was refined with a custom variation of the method to reconcile ANTs-derived and FreeSurfer-derived segmentations of the cortical gray-matter of Mindboggle (Klein et al. 2017). Volume-based spatial normalization to the ICBM 152 Nonlinear Asymmetrical template version 2009c standard space was performed through nonlinear registration with ‘antsRegistration’ from ANTs 2.3.3, using brain-extracted versions of both T1w reference and the T1w template.

BOLD functional data across all sessions and runs were preprocessed collectively. A reference volume and its skull-stripped version were generated using a custom methodology of fMRIPrep. Head-motion parameters with respect to the BOLD reference were estimated, followed by spatiotemporal filtering using ‘mcflirt’ from FSL (Jenkinson et al. 2002). BOLD runs were slice-time corrected to 0.972s (0.5 of slice acquisition range 0s-1.94s) using ‘3dTshift’ from AFNI (Cox and Hyde 1997). The BOLD time-series were resampled onto their original, native space by applying the transforms to correct for head-motion. The BOLD reference was then co-registered to the T1w reference using boundary-based registration via ‘bbregister’ from FreeSurfer (Greve and Fischl 2009). Co-registration was configured with six degrees of freedom. Confounding time-series calculated based on the preprocessed BOLD included: root mean square displacement (RMSD) between frames (Jenkinson et al. 2002), absolute sum of relative framewise displacement (FD) (Power et al. 2014), and the derivative of root mean square variance over voxels (DVARS) (Power et al. 2014), as well as global signals extracted within CSF, WM, and the whole-brain mask. Additionally, a set of physiological regressors were extracted to allow for component-based noise correction (CompCor) (Behzadi et al. 2007). Principal components were estimated after high-pass filtering the preprocessed BOLD time-series (using a discrete cosine filter with 128s cut-off) for the two CompCor variants: temporal (tCompCor) and anatomical (aCompCor). tCompCor components were calculated from the top 2% variable voxels within the brain mask. For aCompCor, three probabilistic masks (CSF, WM, and combined CSF+WM) were generated in anatomical space. For each CompCor decomposition, the k components with the largest singular values that cumulatively explained at least 50% of variance across the nuisance mask (CSF, WM, combined, or temporal) were retained. The BOLD time-series were resampled into standard space with a spatial resolution of 2×2×2 mm^3^ or 97×115×97 voxels. First, a reference volume and its skull-stripped version were generated using a custom methodology of fMRIPrep. All resamplings can be performed with a single interpolation step by composing all the pertinent transformations (i.e., head-motion transform matrices, susceptibility distortion correction when available, and co-registrations to anatomical and output spaces). Gridded (volumetric) resamplings were performed using ‘antsApplyTransforms’ (ANTs), configured with Lanczos interpolation to minimize the smoothing effects of other kernels (Lanczos 1964). Non-gridded (surface) resamplings were performed using ‘mri_vol2surf’ (FreeSurfer).

### First-level modeling

We estimated neural activity in GM voxels for each event of scene or object presentation by constructing first-level general linear models following the *Least Squares – Separate* approach (Mumford et al. 2012), using SPM12 (Friston et al. 2006) and custom MATLAB scripts. Participant-specific GM masks were created by binarizing the fMRIPrep-generated probabilistic masks with a probability threshold of 20%. Each model included one regressor for an event of interest, which was the presentation of either a scene (3 s) or an object (4 s) and another regressor for all other scenes and objects. These two regressors were convolved with the canonical double-Gamma hemodynamic response function, and their temporal and dispersion derivatives were included to allow for variations in the exact timing of peak response. Covariates of no interest included global signal, WM signal, CSF signal, framewise displacement, DVARS, RMSD, and six translational and rotational motion parameters. A high-pass temporal filter of 128 s was applied, and the AR(1) model was used to remove autocorrelation. These first-level models yielded regression coefficients (betas) that estimated voxel-level neural activity corresponding to a specific event.

### Analysis plan

#### Quality checks

First, we examined whether our manipulation of schema congruency achieved its intended effects by plotting histograms of YAs’ and OAs’ ratings on the likelihood of finding the object in the scene. Overall, congruent and incongruent pairs received many ratings on the upper and lower ends of the scale, respectively; however, opinions widely diverged regarding the neutral pairs (see **Figure S1**). Given the lack of consensus, as well as to accommodate individual differences in life experiences and background knowledge, we used each participant’s own ratings to categorize the trials as: [4] = “schema-congruent”, [1] = “schema-incongruent”, and [2,3] = “schema-neutral”; trials that did not receive a rating (3.1%) were excluded from subsequent analyses. The use of subjective judgment of congruency is especially important in this study since the two age groups studied here appeared to disagree on certain pairs (see **Figure S2**).

Then, we examined the subsequent recognition of the objects. Participants apparently varied in their ability to discriminate between old and new objects, and in their decision criterion (see **Figure 2A**), rendering direct comparisons of raw responses difficult. To mitigate such biases, we conducted a receiver-operating characteristics (ROC) analysis using ‘yardstick 1.3.1’ (Kuhn et al. 2024) in R (4.3.3). Two YAs and two OAs with an AUC lower than 0.6 were deemed as low adherence to task and excluded from all subsequent analyses. For the remaining participants (32 YAs and 28 OAs), it was decided whether counting only “4” responses, or both “3” and “4” responses as “old” resulted in a better decision outcome (i.e., closer to 100% true positive and 0% false positive), which was then used to compute the sensitivity index (d’) of recognition in each subjective congruency condition.

**Figure 2.**
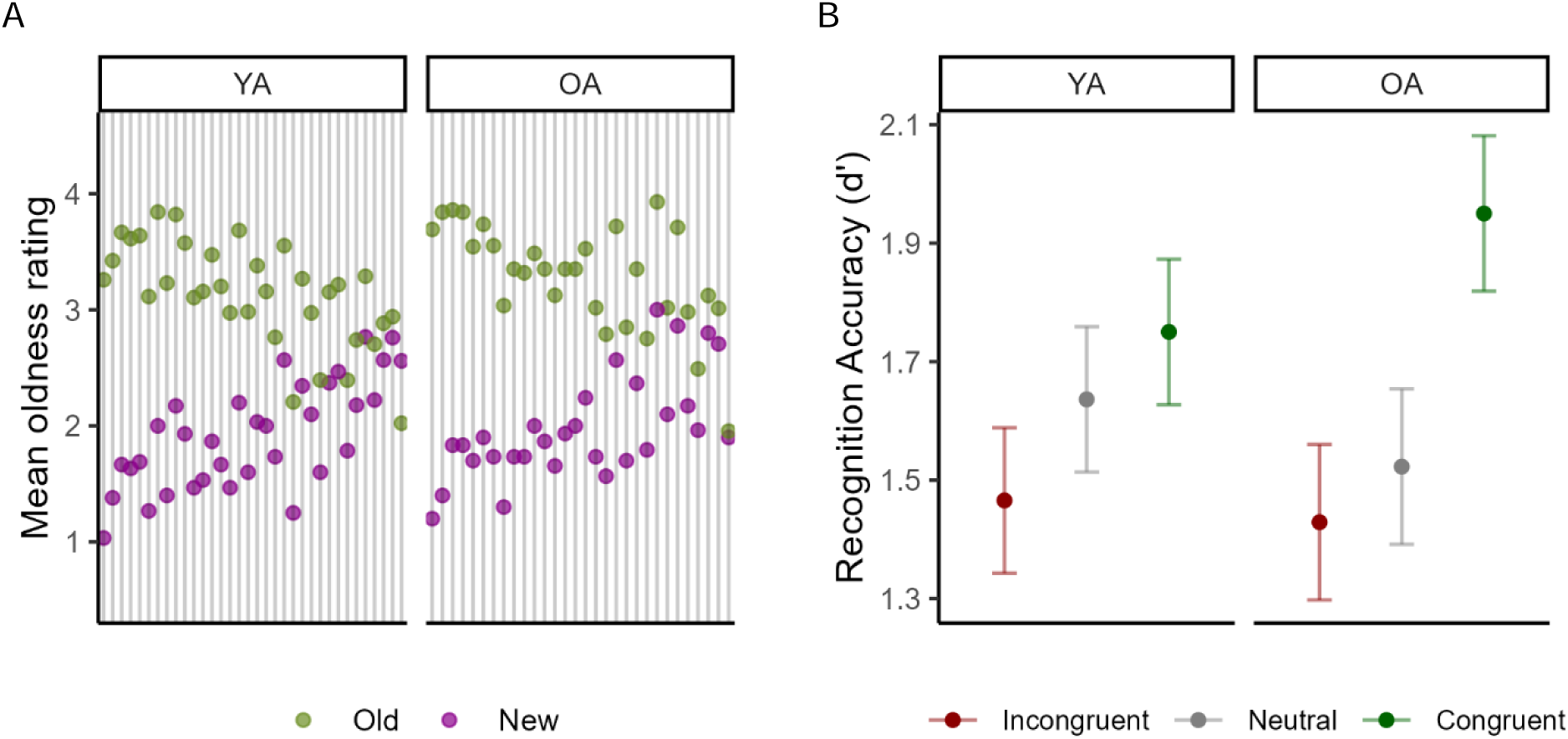
Memory of objects. **A)** Mean oldness ratings (1=“definitely new”, 2=“probably new”, 3=“probably old”, 4=“definitely old”) of old and new objects during the conceptual memory test for each participant. **B)** Recognition accuracy (d’) by age group and congruency. Error bars indicate standard errors. *YA*, younger adults; *OA*, older adults.

#### Congruency and age effects on recognition

To understand how aging and schema congruency may impact object recognition memory, we fitted a linear mixed-effects model that predicted recognition accuracy (d’) with fixed effects of subjective *Congruency* (3 levels: congruent, neutral, and incongruent), *Age Group* (2 levels: YA and OA), and their interaction, using ‘lme4 1.1-35.3’ (Bates et al. 2015) in R. Participants were modeled as random intercepts. P-values were estimated using Satterthwaites’s method via the ‘lmerTest 3.1-3’ package (Kuznetsova et al. 2017). Post-hoc tests of contrasts and interactions were examined using ‘emmeans 1.10.1’ with p-values adjusted for multiple comparisons using the multivariate t-distribution (Lenth 2020).

#### Congruency and subsequent memory effects in cortico-hippocampal interaction

We then examined how object encoding activity in HPC was modulated by schema and semantic processing during scene viewing in the following broad regions: vmPFC, AG, and ATL. Regions of interest were demarcated based on the Human Brainnetome atlas (Fan et al. 2016), with four hippocampal subregions (left anterior, left posterior, right anterior, and right posterior) examined separately (see **Table S1** for coordinates). To test cortico-hippocampal modulations, we used a modified version of psychophysiological interaction (PPI) (Friston et al. 1997; Huang, De Brigard, et al. 2024) analysis using trial-level activity estimates and linear mixed-effects models. First, trial-level activity series for each region was obtained by averaging voxel-wise trial-level beta series. The effect of head motion (FD) was removed from activity series for each brain region via a set of regressions, and trials containing timepoints with large motions (FD > 1 mm) were excluded from subsequent analysis to minimize artifacts. The activity series were also z-scored within participant, congruency level, and subsequent memory to focus on congruency-related subsequent memory effect within each participant. Second, each model predicted object encoding activity of in a HPC subregion with fixed effects of scene viewing activity in a cortical region (e.g., vmPFC), *Congruency* (3 levels: Congruent, Neutral, and Incongruent), *Age Group* (2 levels: YA and OA), and *Subsequent Memory* (2 levels: Remembered and Forgotten). Random effects included random intercepts of object identities and random slopes of cortical scene viewing activity by participant. This modified PPI model effectively estimated how much a cortical region modulated hippocampal encoding process in either age group, given different degrees of schematic information, and in relation to subsequent memory.

Notably, our analysis differed from the conventional PPI analysis in two key aspects. First, instead of BOLD signal time courses, our “physiological” variables were neural activity series estimated at the level of experimental trials, i.e., a scene or an object (Rissman et al. 2004; Mumford et al. 2012). This way, statistical interactions were tested at the neural level, which could be more informative of the underlying cognitive processes than interactions at the hemodynamic level (Di et al. 2017). Second, instead of taking the usual two-step procedure that first fits separate models for individuals and then performs group-level tests of participant-level estimates, we chose a mixed-effects model approach, which offers better sensitivity and generalizability (Baayen et al. 2008; Brauer and Curtin 2018). Moreover, because both Congruency and Subsequent Memory were individually determined by participant behavior during the experiment, the number of trials across cells (e.g., “participant A & Congruent & Forgotten”) was inevitably unequal. While the conventional two-step PPI does not account for such an imbalance, mixed-effects models do so through partial pooling (Chen et al. 2021) and are thus much more appropriate for the present study. Importantly, the focal outcome from these models was the subsequent memory effect (“Remembered - Forgotten” contrast) in the regression coefficients of scene viewing activity, which indicates the mnemonic relevance of the assessed cortico-hippocampal modulation. P-values were adjusted for false discovery rate (FDR) across four HPC subregions, and significance was determined using a threshold of q < 0.05.

#### Individual differences in schematic modulation and semantic representation

To test our hypothesis that the observed AG-HPC interactions reflected semantic-memory-mediated episodic memory encoding, we examined left AG’s representational strength of scene semantic information using representational similarity analysis (Kriegeskorte et al. 2008). To this end, we converted the labels of each scene into a vector of length 300 using a pre-trained word2vec model (word2vec 2013), based on which we computed the pairwise cosine similarity across all scenes and obtained a 114-by-114 semantic similarity matrix. Then, we computed all pairwise Pearson’s correlation of left AG activity patterns of scenes, obtaining a 114-by-114 neural pattern similarity matrix. Finally, the vectorized lower triangular parts of the two similarity matrices were compared using Spearman’s rank correlation. The Fisher’s z-transformed correlation coefficient indexed left AG’s representational strength of scene meaning.

Furthermore, we extracted from the fitted mixed-effects model the random slopes, which indexed individual deviations in the strength of cortico-hippocampal modulations. We examined the correlation between AG-HPC random slopes and left AG semantic representation strength, and between AG-HPC random slopes and vmPFC-HPC random slopes. Correlation coefficients for YA and OA groups were compared using ‘cocor 1.1-4’ (Diedenhofen and Musch 2015).

## Results

### Congruency effects on memory by age groups

Behavioral results are shown in **Figure 2** and **Table S2**. We first assessed participants’ memory of objects by testing the recognition accuracy (d’). We found no significant main effect of *Age Group* (*F*_1, 58_ = 0.01, p = 0.9235) but a significant main effect of *Congruency* (*F*_2, 116_ = 28.53, p < 0.0001), as well as a significant *Congruency* × *Age Group* interaction (*F*_2, 116_ = 4.52, p = 0.0129). Post-hoc tests examining the age groups separately revealed that, in YAs, schema-congruent objects (mean_con_ = 1.75) were better remembered than schema-incongruent objects (mean_inc_ = 1.47; b = 0.28, SE = 0.07, t_116_ = 3.83, p = 0.0012) but not better than schema-neutral ones (mean_neu_ = 1.64; b = 0.11, SE = 0.07, t_116_ = 1.53, p = 0.4791); however, in OAs, schema-congruent objects (mean_con_ = 1.95) were better remembered than both schema-incongruent (mean_inc_ = 1.43; b = 0.52, SE = 0.08, t_116_ = 6.56, p < 0.0001) and schema-neutral objects (mean_neu_ = 1.52; b = 0.43, SE = 0.08, t_116_ = 5.38, p < 0.0001). Moreover, OAs showed a larger congruency-related memory improvement than YAs (b = 0.31, SE = 0.11, t_116_ = 2.88, p = 0.0129). These results remain the same after statistically controlling for NIH crystallized intelligence scores, thereby excluding the possibility that the differential recognition accuracy was simply due to an age-related difference in general knowledge (see **Table S3**). In sum, the behavioral result confirms our prediction of a larger congruency effect in OAs than YAs, consistent with the idea that OAs over-rely on semantic schemas, boosting subsequent memory for schema-congruent information.

### VmPFC’s and AG’s modulations of HPC encoding process

To understand the neural mechanisms underlying how available schemas and semantic knowledge facilitate episodic memory encoding in YAs and OAs, we assessed the subsequent memory effect in the interplay between schema-related regions (vmPFC, AG, and ATL) and HPC. Below the results for vmPFC and AG are reported. For ATL, no cortico-hippocampal modulation effect was significant (p_FDR_ > 0.11; see **Table S4**).

**vmPFC**. We found a significant main effect of *Congruency* on vmPFC’s modulation of left anterior HPC (**Figure 3A**, *F*_2, 6240_ = 6.74, p = 0.0012, p_FDR_ = 0.0036) and left posterior HPC (**Figure 3B**, *F*_2, 6258_ = 5.60, p = 0.0037, p_FDR_ = 0.0112), with no significant main effect of *Age Group* or interaction (p > 0.72). Post-hoc tests across age groups revealed a significant subsequent memory effect (i.e., “Remembered > Forgotten”) for schema-congruent objects (anterior: b = 0.19, SE = 0.07, t_6247_ = 2.92, p = 0.0035; posterior: b = 0.21, SE = 0.07, t_6247_ = 3.06, p = 0.0022) but not for schema-neutral ones (p > 0.52). Moreover, vmPFC’s modulation of left anterior HPC showed a significant subsequent forgetting effect (i.e., “Remembered < Forgotten”; b = -0.09, SE = 0.05, t_6226_ = -2.09, p = 0.0363). These results suggest that, for both YAs and OAs, accessing schematic knowledge about scenes via vmPFC activation positively facilitated left HPC encoding of objects that are congruent with the schema, while activation of schemas also impeded the encoding of incongruent and unexpected objects.

**Figure 3.**
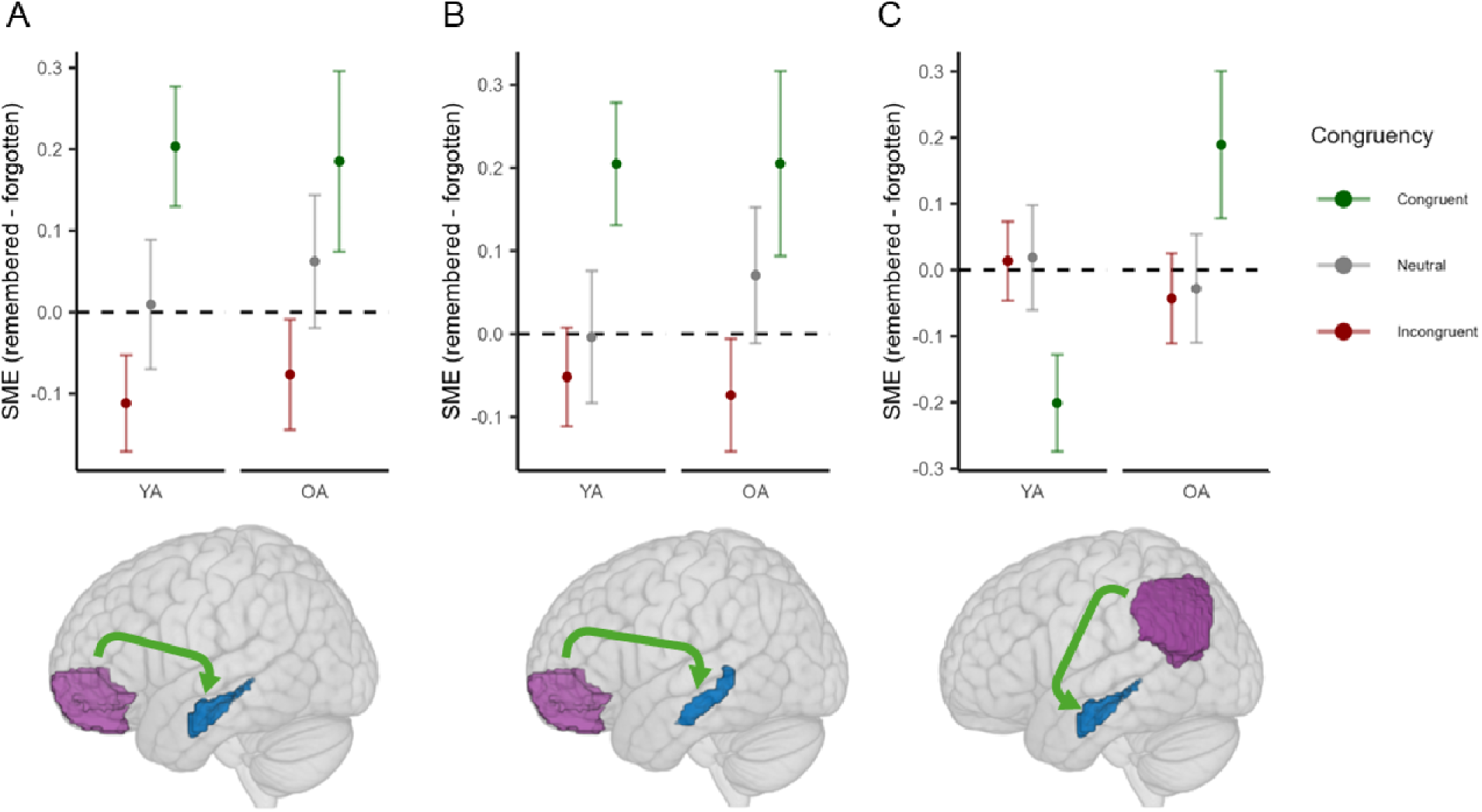
Cortico-hippocampal modulations. Subsequent memory effects (SME) in different cortical modulations of hippocampal encoding processes by Age Group and Congruency. **A**) vmPFC’s modulation of left anterior HPC. **B**) vmPFC’s modulation of left posterior HPC. **C**) Left AG’s modulation of left anterior HPC. Error bars indicate standard errors. *vmPFC*, ventromedial prefrontal cortex; *AG*, angular gyrus; *HPC*, hippocampus.

**AG**. With respect to left AG’s modulation of left anterior HPC, we found a significant *Congruency* × *Age Group* interaction effect on left AG’s modulation of left anterior HPC (*F*_2, 6254_ = 4.38, p = 0.0126, p_FDR_ = 0.0379). Post-hoc tests revealed that there was no age-related difference for objects that were unexpected (incongruent: b = - 0.06, SE = 0.09, t_6169_ = -0.63, p = 0.5319) or unrelated to the schema (neutral: b = -0.05, SE = 0.11, t_6253_ = -0.41, p = 0.6831). However, for schema-congruent objects, the subsequent memory effect in the modulation was stronger in OAs than in YAs (b_OA_ = 0.19, b_YA_ = -0.20; b_OA-YA_ = 0.39, SE = 0.13, t_6229_ = 2.93, p = 0.0034; see **Figure 3**). Given left AG’s role in semantic memory, this result suggests that OAs might benefit from their rich general knowledge about the scene to facilitate later encoding of the object, provided that they were semantically associated.

### Individual differences in cortico-hippocampal modulations

To further examine the purported benefit of semantic knowledge specific to OAs, we related individual differences in left AG’s modulation of left anterior HPC to the representational strength of semantic information of scenes in left AG. We found that, although the difference in correlation coefficients between age groups was not significant (z = -1.59, p = 0.1126), modulatory strength and semantic representational strength in left AG were positively correlated in OAs (r = 0.45, t_26_ = 2.58, p = 0.0160) but not in YAs (r = 0.05, t_30_ = 0.29, p = 0.7738; see **Figure 4A**).

**Figure 4.**
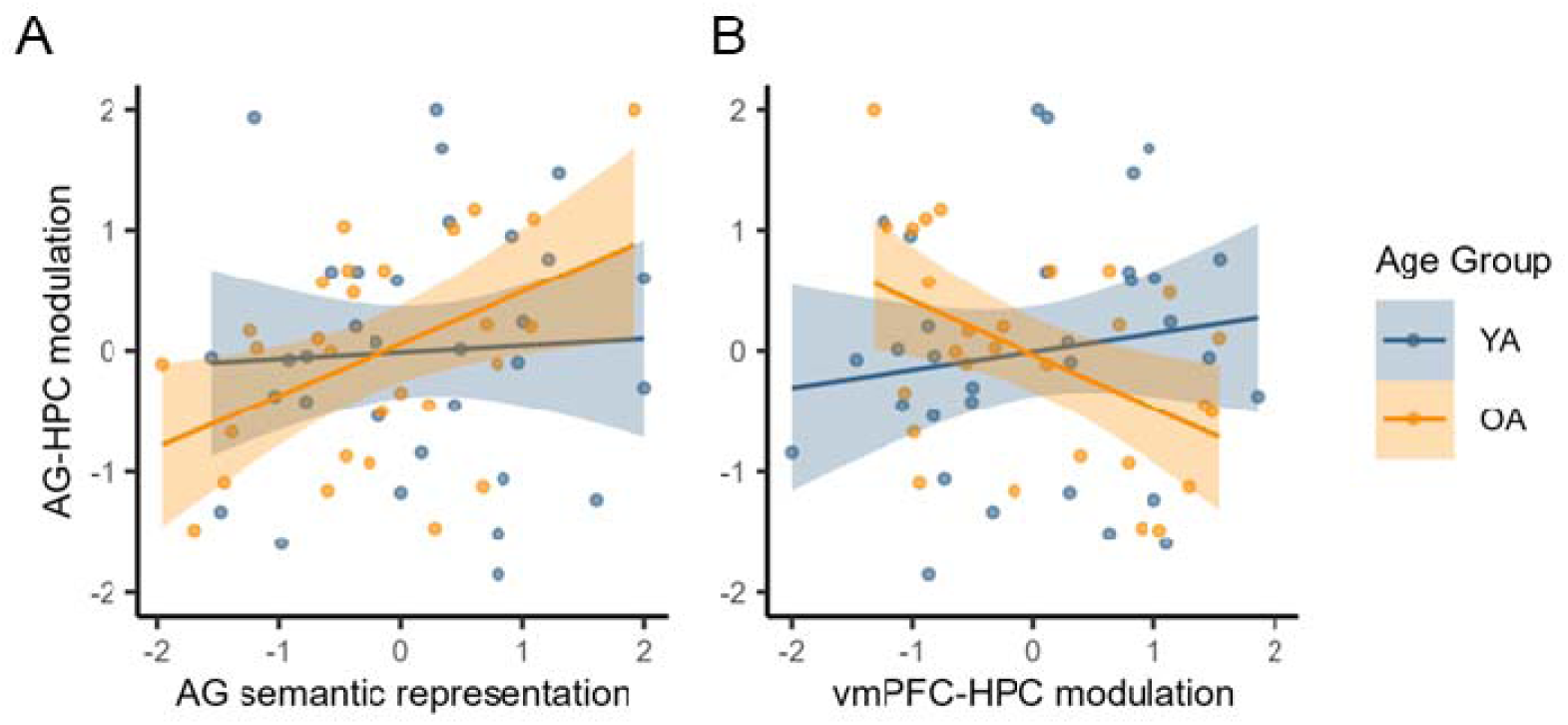
Individual differences. Individual differences in the left angular gyrus’s (AG’s) modulation of left anterior hippocampus (HPC) against **A)** representational strength of scene semantic information in left AG and **B)** individual differences in ventromedial prefrontal cortex’s (vmPFC’s) modulation of left anterior HPC. Values were z-scored and outliers (|*z*|>2) were clipped.

Furthermore, we assessed the relationship between the strength of modulations from vmPFC and left AG. For OAs, a positive across-participant correlation of modulatory strengths would suggest the presence and benefit of a well-integrated schema-semantics-encoding network or process-specific alliance (Cabeza et al., 2018) that have potentially arisen from years of life experiences, whereas a negative correlation would suggest that OAs use semantic knowledge to facilitate episodic memory encoding as a compensatory strategy. Comparing random slopes from the above fitted models, we found no significant correlation in YAs (r = 0.15, t_30_ = 0.81, p = 0.4262), but a significant negative correlation in OAs (r = -0.47, t_26_ = -2.73, p = 0.0112). The difference in correlations between OAs and YAs was also significant (z = 2.42, p = 0.0156; see **Figure 4B**), consistent with the second hypothesis: OAs exploited their rich semantic knowledge of the scenes to complement their encoding of object concepts.

## Discussion

In the current fMRI study, we focused on an important open question in the cognitive neuroscience of aging, namely, how OAs take advantage of their rich knowledge for semantic schemas to boost their encoding of new information. We focused on congruency effects in scene-object paradigm, and on cortical-HPC modulation predicting subsequent memory. We report four main findings: (1) congruency enhanced subsequent episodic memory to a greater extent in OAs than in YAs; (2) HPC contribution to subsequent memory was modulated by vmPFC similarly in both age groups; (3) HPC contribution to subsequent memory was modulated by left AG in OAs but not in YAs; and (4) finally, in OAs, the modulatory effects of AG was positively correlated with its semantic representation strength and negatively correlated with the modulatory effects of vmPFC. These results support the idea that OAs counteract episodic memory deficits by relying on semantic schemas, and identify AG modulation as a potentially complementary mechanism for the vmPFC mechanism shared with YAs. We discuss the four main findings in separate sections below.

### Schema congruency enhanced memory to a greater extent in OAs than YAs

Our results clearly show that congruency with schemas enhanced memories, and that this enhancement was more prominent in OAs than in YAs. The general beneficial effect of schema congruency is consistent with multiple findings in the literature (e.g., Brewer and Treyens 1981; De Brigard et al. 2017; van Kesteren et al. 2020). Congruency with available schemas is theorized to facilitate the encoding process, since schemas provide a knowledge scaffold unto which new congruent information can be efficiently assimilated (Ghosh and Gilboa 2014; Umanath and Marsh 2014). Schemas also enable inferential elaboration of new information and may lead to stronger memory traces (Anderson 1984; Ghosh and Gilboa 2014).

Compared to neutral pairs, congruent pairs involved not only a better fit with scene schemas (i.e., objects typically included in a particular scene) but also stronger semantic associations between scene and object names (e.g., *garden-rose* vs. *garden-television*). Thus, better memory in the congruent condition could also be in part due to semantic relatedness (Bein et al. 2015). In fact, OAs are impaired when learning semantically unrelated word pairs, but they can perform as well as YAs when learning semantically related word pairs (Delhaye et al. 2019; Bi 2021). It is an open question for future research whether the effects of schematic support in OAs involve the same or different mechanisms as the general effects of semantic support.

Critically, our behavioral results confirmed our prediction of a stronger congruency effect of memory in OAs than in YAs. This result is consistent with prior studies (Smith et al. 1998; Chen et al. 2022), and supports the idea that OAs utilize preserved semantic memory to facilitate their likely declined episodic memory (Umanath and Marsh 2014). Our neuroimaging findings help to outline the potential neural mechanisms underlying this behavioral effect.

### Hippocampal contribution to subsequent memory was modulated by vmPFC in both age groups

The main goal of this fMRI study was to examine how schematic information processing in vmPFC, AG, and ATL modulates the success of object memory encoding in HPC, and how such cortico-hippocampal interactions operate at distinct levels of schema congruency and in different age groups. Intuitively, the more congruent an object is with the currently instantiated schema (scene), the more schematic information can facilitate the encoding and improve the subsequent memory of the object. This idea is supported by our finding that, in both age groups, the impact of the left HPC on subsequent memory was modulated by vmPFC in a congruency-related manner, such that the degree of modulation increased with schema congruency. This finding is consistent with past work showing that, given schematic knowledge, stronger vmPFC-HPC functional connectivity is associated with better subsequent memory across participants (Liu et al. 2017).

It has been suggested that the medial PFC directs the assimilation of congruent information, whereas the medial temporal lobe mediates the integration of novel and arbitrary information (van Kesteren et al. 2012; van Kesteren et al. 2013). Further, it has been hypothesized that medial PFC actively inhibits the medial temporal lobe. Consistent with this hypothesis, there is evidence of *reduced* frontal-temporal functional connectivity in the presence of schematic knowledge (van Kesteren et al. 2010; Bein et al. 2014). The difference between these findings and our finding of stronger vmPFC-HPC connectivity for congruent pairs most likely reflect differences in the paradigms employed. Unlike many past studies, in the current study, scene and object stimuli were presented not *simultaneously* but *serially*. This feature also allowed us to model the presentation of each scene and object in the pair as separate events with their own neural activity, instead of treating them as a single, undifferentiated event (cf. van Kesteren et al. 2013).

Finally, it is interesting to note that although we did not observe an age-related difference in vmPFC-HPC interaction, there is evidence that, during successful memory encoding, OAs sometimes show greater vmPFC-HPC connectivity than YAs (Dennis et al. 2008; Addis et al. 2010). A possible explanation is that OAs may have a tendency to rely on vmPFC-mediated schematic knowledge to support novel episodic encoding, but YAs do the same when schematic knowledge is made explicit as in this current study.

### Hippocampal contribution to subsequent memory was modulated by left AG in OAs

We also examined how schematic knowledge processing in left AG modulated object memory encoding in HPC. These two regions are strongly associated by extensive structural and functional connections (Uddin et al. 2010; Hermiller et al. 2019; Huang, Howard, et al. 2024). Also, AG is thought to facilitate schema-related processes (Gilboa and Marlatte 2017; van der Linden et al. 2017), given its role in binding multisensory information (Bonner et al. 2013; Bonnici et al. 2016; Humphreys et al. 2021) and semantic processing (Price et al. 2015; Kuhnke et al. 2023). Moreover, one study found that AG-HPC functional connectivity tracked schema congruency (van der Linden et al. 2017). However, what was not known was the impact of this connection on subsequent memory in YAs and OAs.

The current results showed that left AG modulated left anterior HPC to facilitate subsequent memory. Most interestingly, we observed a significant *Congruency* × *Age Group* interaction, which clearly shows that OAs exhibited a greater improvement in subsequent memory than YAs due to this AG-HPC interaction. This finding suggests that OAs preferentially make use of their rich semantic knowledge of scenes to facilitate episodic memory encoding, consistent with the schematic support effect in cognitive aging (Umanath and Marsh 2014).

### Negative correlation between vmPFC and AG effects in OAs

To further validate the schematic support interpretation of this differential AG-HPC interaction between age groups, we examined the representation of scene semantics in left AG. In OAs, the strength of semantic representation in left AG was positively correlated with the strength of its modulation of HPC, whereas these two measures were not correlated in YAs. Additionally, the modulatory strengths from left AG and from vmPFC were negatively correlated in OAs but not in YAs. These two results collectively support the idea that OAs exploit their semantic memory to facilitate the encoding of schema-congruent objects as a compensatory mechanism for their declined episodic memory.

### Open questions

The current results open several questions for future studies, and here we highlight two of them. One interesting question is the particular HPC subregion where memory processes are modulated by schemas in YAs and OAs. Consistent with past research, in the current study, the schematic modulations of HPC by both the vmPFC and left AG were found within the *left anterior* HPC. Past research examining the functional distinction of hippocampal subregions between hemispheres (Ezzati et al. 2016), along the long axis (Sekeres et al. 2018; Grady 2020; Gardette et al. 2022), or both (Persson and Söderlund 2015), suggests that left and/or anterior HPC specifically supports the coding of coarse-grained gist information (as opposed to fine-grained details) and is preferentially recruited during the encoding of verbal (as opposed to pictorial) materials. In the current study, participants were shown both visual images and verbal labels of scenes and objects, but memory of objects was cued with verbal labels. This experimental design led to the *conceptual* information of objects being highly relevant and appropriate for subsequent memory, according to the transfer-appropriate processing account (Morris et al. 1977; Bramão and Johansson 2018; Huang, Howard, et al. 2024), which is consistent with the fact that our findings of cortical modulations especially acted on left anterior HPC. Further research could investigate the role of schemas for conceptual and perceptual forms of memory.

Another interesting question for future studies is whether the neural mechanisms of congruency and incongruency effects are expressed similarly across the lifespan, and what design modifications can highlight or suppress these differences. As noted in the Introduction, improved memory for incongruent objects in the scene-object paradigm is typically found when the objects are embedded in the scene (e.g., a chainsaw on top of a kitchen counter) (Mäntylä and Bäckman 1992; Wynn et al. 2020; Klever et al. 2023), whereas the surprise and the incongruency effect largely diminish when objects are separated from the scene (Brod and Shing 2019; Cook et al. 2021). In the current study, we focused on the congruency effect and the schematic support effect in OAs, using an experimental design that separated the scene and the object in both space and time – nonoptimal for eliciting incongruency effects. Given pronounced age-related changes in the structure and function of the HPC (Thomann et al. 2013; Nyberg 2017; Lai and Chang 2023), it is reasonable to speculate that our design promoted active binding strategies that are reliant on hippocampal interactions. In contrast, designs that emphasize incongruencies in the context of rich schematic environments, when older adults are expecting more familiar schematic relationships (Ramey et al. 2024), may instead emphasize age-related changes in monitoring in regions like the vmPFC (Lighthall et al. 2014). Future studies may therefore explore such alternative experimental designs to focus on the neural mechanisms underlying the incongruency effect and examine how they vary by age group.

## Conclusion

In the current fMRI study, we investigated behavioral and neural differences in the schematic support effect on episodic memory in YAs and OAs. Consistent with our hypotheses, we found that OAs demonstrated a stronger schematic support effect than YAs. VmPFC modulated the encoding process in HPC commonly in both age groups. Additionally, left AG in OAs additionally modulated HPC to support subsequent memory, which serves as a compensatory mechanism for deficient vmPFC-HPC connections. Collectively, our findings illustrate age-related differences in how schemas influence episodic memory encoding via distinct routes of cortico-hippocampal interactions.

## CRediT Author contributions

- Shenyang Huang (Investigation, Data Curation, Formal Analysis, Visualization, Writing – Original Draft, Writing – Review & Editing),
- Paul C. Bogdan (Data Curation, Writing – Review & Editing),
- Cortney M. Howard (Data Curation, Writing – Review & Editing),
- Kirsten Gillette (Investigation, Data Curation, Writing – Review & Editing),
- Lifu Deng (Conceptualization, Methodology, Investigation, Data Curation, Writing – Review & Editing),
- Erin Welch (Methodology, Investigation, Data Curation, Writing – Review & Editing),
- Margaret McAllister (Investigation, Data Curation, Writing – Review & Editing),
- Kelly S. Giovanello (Funding Acquisition, Supervision, Writing – Review & Editing),
- Simon W. Davis (Conceptualization, Methodology, Funding Acquisition, Supervision, Writing – Original Draft, Writing – Review & Editing), and
- Roberto Cabeza (Conceptualization, Methodology, Funding Acquisition, Supervision, Writing – Original Draft, Writing – Review & Editing)

## Funding

This work was supported by the National Institutes of Health R01-AG066901.

## Conflict of Interest

All authors confirm that there is no conflict of interest.

## Notes

### Competing Interest Statement

The authors have declared no competing interest.

