## Supplementary Information for "Cortico-hippocampal interactions underlie schema-supported memory encoding in older adults"

### Figure S1. Subjective ratings of congruency

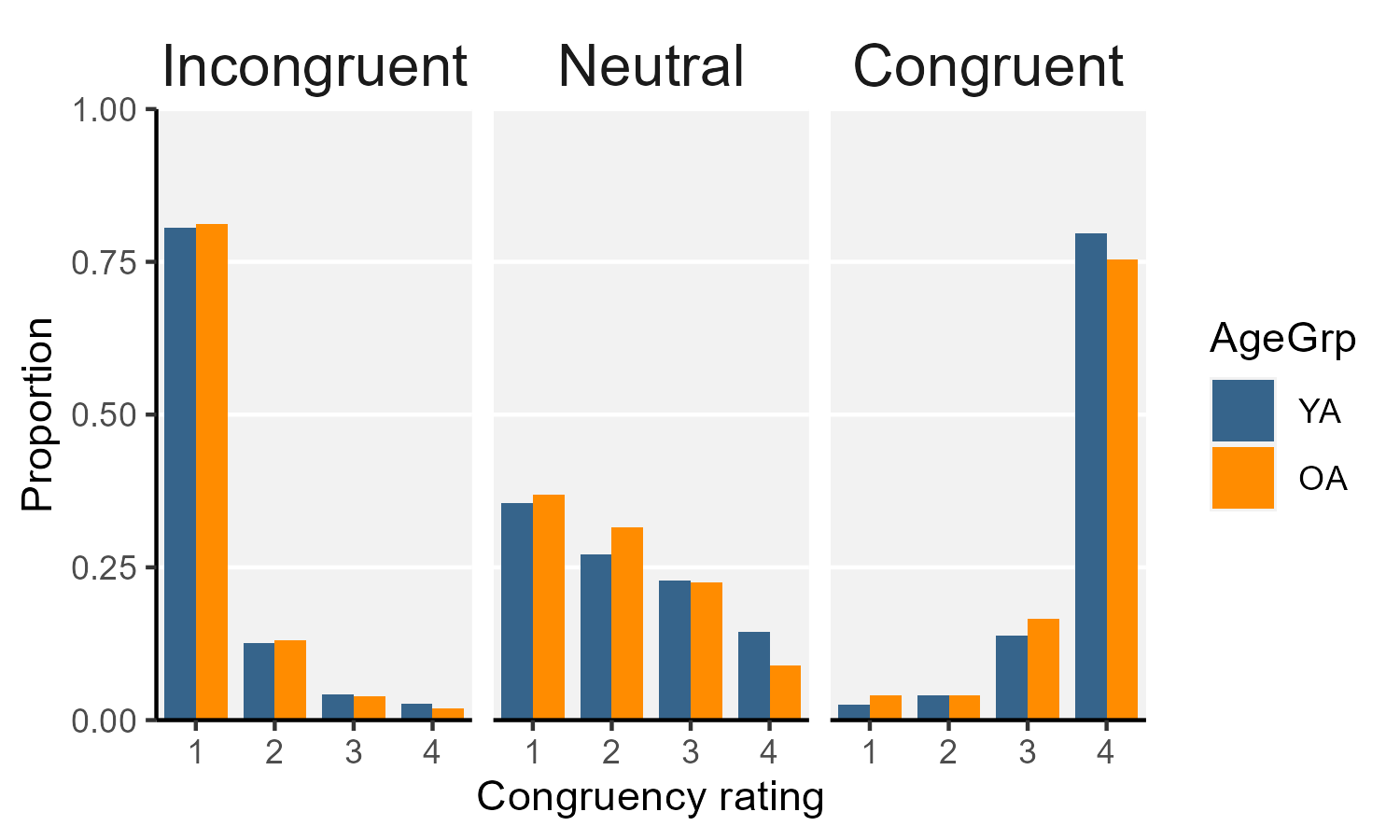

**Figure S1.** Subjective ratings of congruency. The proportion of subjective congruency ratings (1-4) split by *a priori* congruency assignment and age group. Overall, congruent and incongruent pairs received many ratings on the upper and lower ends of the scale, respectively; however, opinions widely diverged regarding the neutral pairs (see **Figure S2**). Therefore, all analyses were conducted using each participant’s own judgment of congruency.

### Figure S2. Subjective ratings of congruency

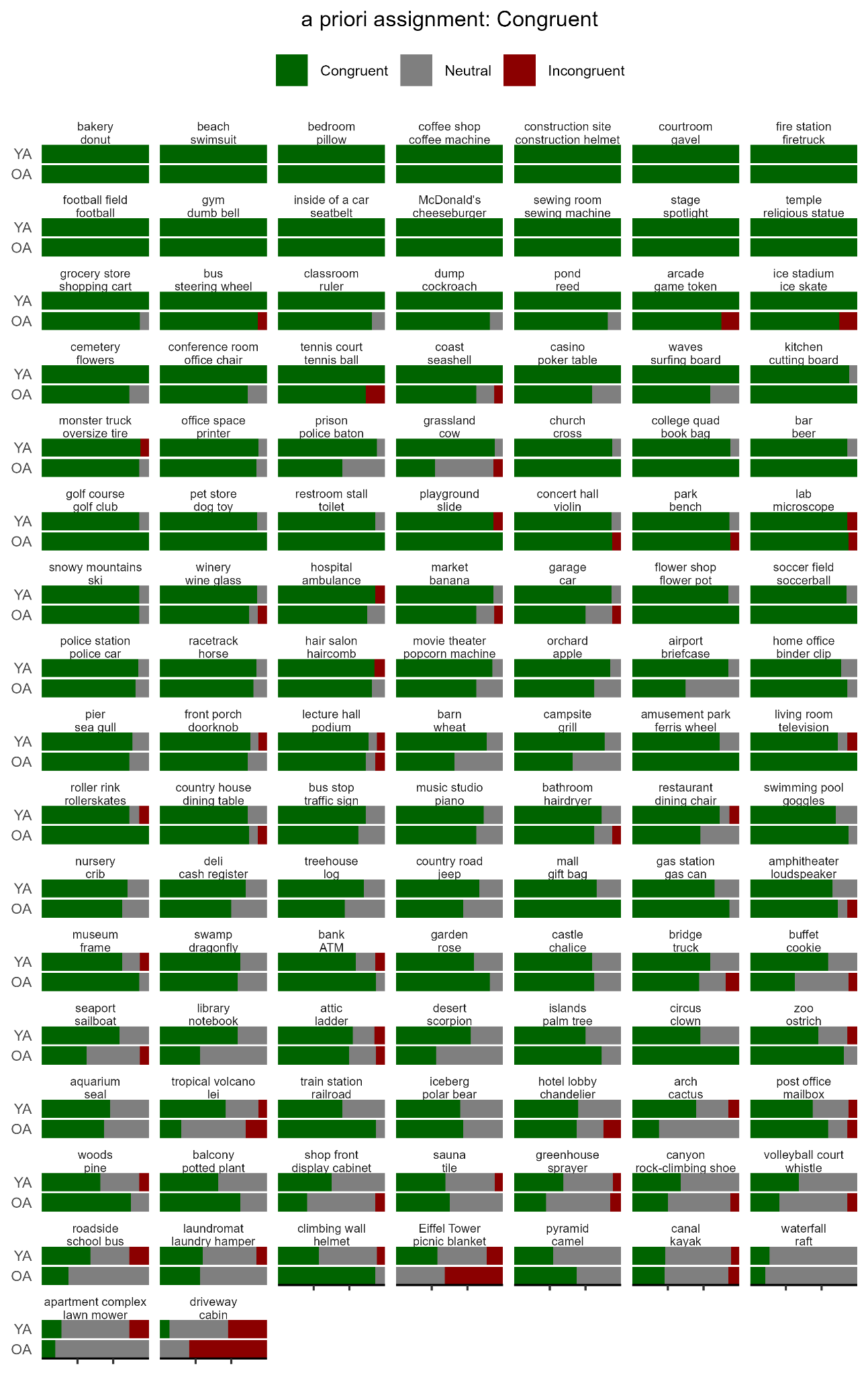

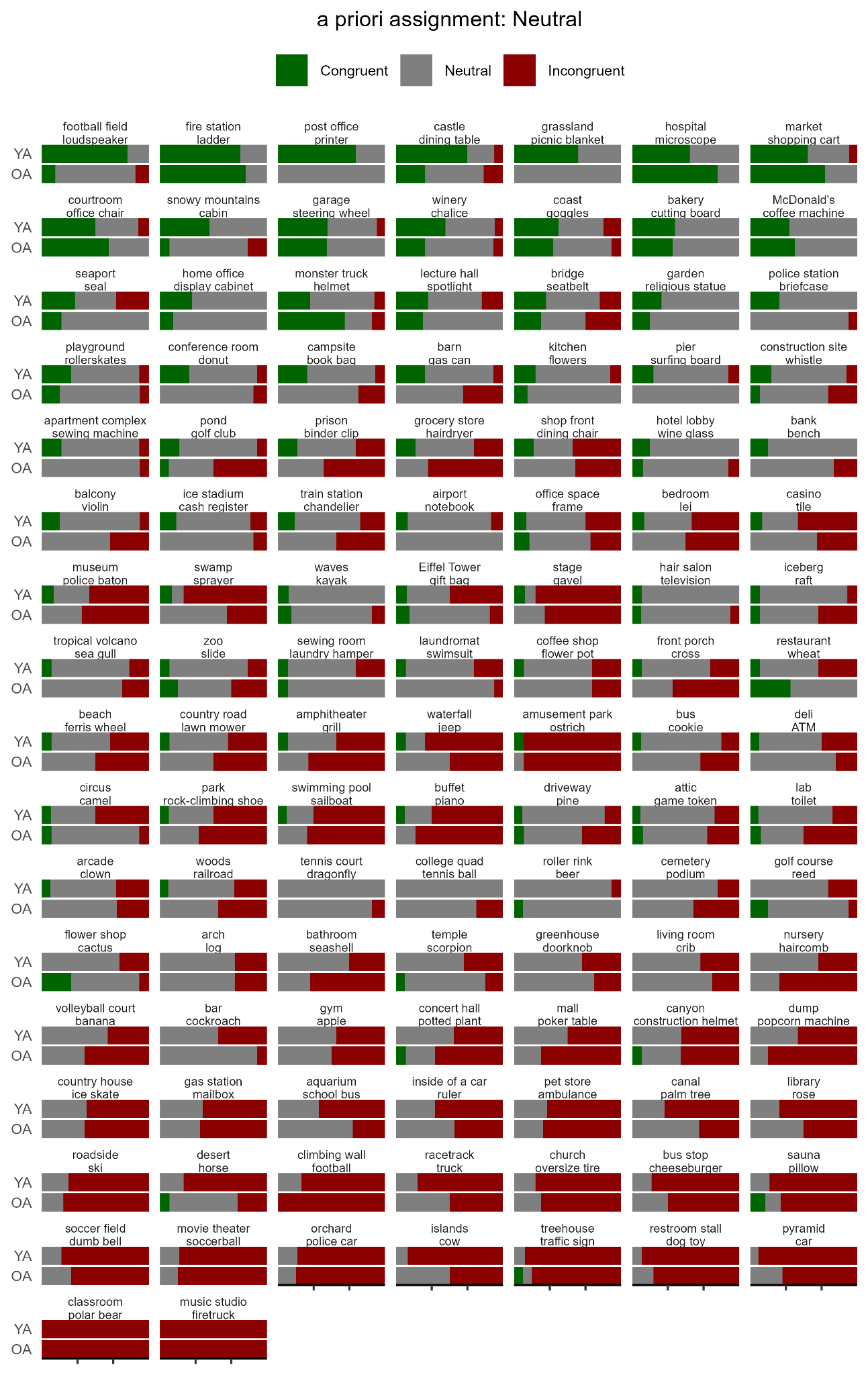

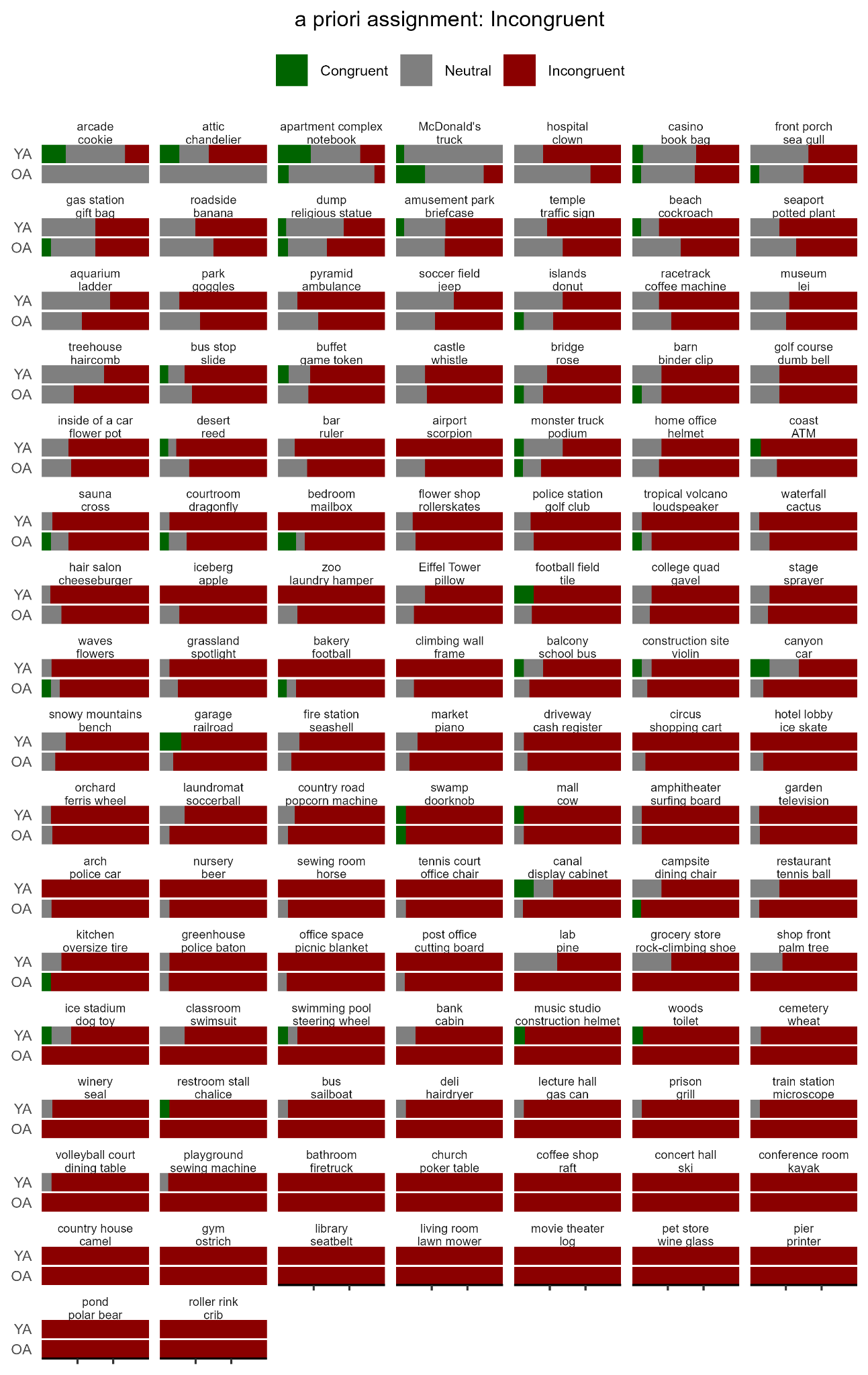

**Figure S2.** Proportion of subjective judgment of congruency ([4] = “congruent”, [1] = “incongruent”, and [2,3] = “neutral”) for each scene-object pair, organized by our *a priori* assignment of congruency. Opinions widely diverged regarding the neutral pairs. Therefore, all analyses were conducted using each participant’s own judgment of congruency.

### Table S1. Regions of interest

| **Name** | **Subregion** | | **MNI coordinates (mm)** | | | **Size (voxel)** |
| --- | --- | --- | --- | --- | --- | --- |
|  |  |  | ***x*** | ***y*** | ***z*** |  |
| vmPFC | - | | 4 | 54 | -11 | 2516 |
| ATL | L | | -41 | 10 | -31 | 1491 |
|  | R | | 46 | 12 | -31 | 1605 |
| AG | L | | -45 | 57 | 36 | 3454 |
|  | R | | 52 | -53 | 38 | 3488 |
| HPC | L | Ant | -21 | -13 | -17 | 560 |
|  |  | Pos | -27 | -29 | -9 | 557 |
|  | R | Ant | 24 | -11 | -19 | 459 |
|  |  | Pos | 32 | -25 | -9 | 614 |

**Note**. Abbreviations: *vmPFC*, ventromedial prefrontal cortex; *ATL*, anterior temporal lobe; *AG*, angular gyrus; *HPC*, hippocampus; *L*, left; *R*, right; *Ant*, anterior; *Pos*, posterior.

| **Table S2. Memory of objects** | | | | | |
| --- | --- | --- | --- | --- | --- |
| **Predictor** | | **b (95% CI)** | **SE** | **t** | **p** |
| Intercept | *-* | 1.64 (1.39 – 1.88) | 0.13 | 13.08 | <0.0001 |
| Age Group | *OA* | -0.11 (-0.48 – 0.25) | 0.18 | -0.62 | 0.5362 |
| Congruency | *Congruent* | 0.11 (-0.03 – 0.26) | 0.07 | 1.53 | 0.1273 |
|  | *Incongruent* | -0.17 (-0.32 – -0.02) | 0.07 | -2.30 | 0.0228 |
| Age Group × Congruency | *OA:Congruent* | 0.31 (0.10 – 0.53) | 0.11 | 2.88 | 0.0044 |
|  | *OA:Incongruent* | 0.08 (-0.14 – 0.29) | 0.11 | 0.71 | 0.4807 |

**Note.** Reference levels: *neutral* (Congruency), *YA* (Age Group). Degrees of freedom = 172.

### Table S3. Memory of objects, controlling for crystallized intelligence

| **Predictor** | | **b (95% CI)** | **SE** | **t** | **p** |
| --- | --- | --- | --- | --- | --- |
| Intercept | *-* | 1.64 (1.39 – 1.88) | 0.12 | 13.33 | <0.0001 |
| Age Group | *OA* | -0.11 (-0.47 – 0.24) | 0.18 | -0.63 | 0.5283 |
| Congruency | *Congruent* | 0.11 (-0.03 – 0.26) | 0.07 | 1.53 | 0.1291 |
|  | *Incongruent* | -0.17 (-0.32 – -0.02) | 0.07 | -2.29 | 0.0234 |
| Age Group ×  Congruency | *OA:Congruent* | 0.31 (0.10 – 0.53) | 0.11 | 2.87 | 0.0046 |
|  | *OA:Incongruent* | 0.08 (-0.14 – 0.29) | 0.11 | 0.70 | 0.4827 |
| Crystallized intelligence | *-* | 0.17 (-0.07 – 0.42) | 0.12 | 1.41 | 0.1608 |
| Age Group ×  Crystallized intelligence | *OA:Cryst* | 0.04 (-0.32 – 0.40) | 0.18 | 0.23 | 0.8219 |
| Congruency ×  Crystallized intelligence | *Congruent:Cryst* | 0.03 (-0.12 – 0.18) | 0.08 | 0.37 | 0.7134 |
|  | *Incongruent:Cryst* | 0.07 (-0.08 – 0.22) | 0.08 | 0.97 | 0.3338 |
| Age Group ×  Congruency ×  Crystallized intelligence | *OA:Congruent:Cryst* | -0.13 (-0.35 – 0.08) | 0.11 | -1.22 | 0.2226 |
|  | *OA:Incongruent:Cryst* | -0.16 (-0.38 – 0.06) | 0.11 | -1.46 | 0.1460 |

**Note**. Reference levels: *neutral* (Congruency),*YA* (Age Group). Denominator degrees of freedom = 166.

### Table S4. Cortico-hippocampal modulations

| **Cortical region** | **HPC subregion** | **Predictor** | **F ratio** | **df** | **p** | **p_FDR_** |
| --- | --- | --- | --- | --- | --- | --- |
| vmPFC | L Ant | Age Group | 0.12 | (1, 6104) | 0.7274 | 0.7274 |
|  |  | Congruency | 6.74 | (2, 6240) | 0.0012 | 0.0036 |
|  |  | Age Group × Congruency | 0.09 | (2, 6245) | 0.9152 | 0.9152 |
|  | L Pos | Age Group | 0.07 | (1, 5861) | 0.7932 | 0.7932 |
|  |  | Congruency | 5.60 | (2, 6258) | 0.0037 | 0.0112 |
|  |  | Age Group × Congruency | 0.23 | (2, 6262) | 0.7970 | 0.7970 |
|  | R Ant | Age Group | 0.05 | (1, 6180) | 0.8168 | 0.8505 |
|  |  | Congruency | 2.40 | (2, 6246) | 0.0912 | 0.2737 |
|  |  | Age Group × Congruency | 0.56 | (2, 6250) | 0.5743 | 0.8614 |
|  | R Pos | Age Group | 0.13 | (1, 6049) | 0.7176 | 0.7795 |
|  |  | Congruency | 0.48 | (2, 6263) | 0.6185 | 0.9277 |
|  |  | Age Group × Congruency | 0.25 | (2, 6266) | 0.7777 | 0.7777 |
| L AG | L Ant | Age Group | 2.05 | (1, 5862) | 0.1520 | 0.2522 |
|  |  | Congruency | 0.01 | (2, 6253) | 0.9882 | 0.9882 |
|  |  | Age Group × Congruency | 4.38 | (2, 6254) | 0.0126 | 0.0379 |
|  | L Pos | Age Group | 0.08 | (1, 5936) | 0.7739 | 0.7932 |
|  |  | Congruency | 0.20 | (2, 6263) | 0.8165 | 0.9672 |
|  |  | Age Group × Congruency | 1.36 | (2, 6264) | 0.2558 | 0.7675 |
| R AG | R Ant | Age Group | 0.04 | (1, 5968) | 0.8505 | 0.8505 |
|  |  | Congruency | 0.19 | (2, 6256) | 0.8251 | 0.8251 |
|  |  | Age Group × Congruency | 2.24 | (2, 6247) | 0.1067 | 0.3200 |
|  | R Pos | Age Group | 0.08 | (1, 5838) | 0.7795 | 0.7795 |
|  |  | Congruency | 0.60 | (2, 6266) | 0.5479 | 0.9277 |
|  |  | Age Group × Congruency | 2.27 | (2, 6258) | 0.1031 | 0.3094 |
| L ATL | L Ant | Age Group | 1.90 | (1, 6127) | 0.1681 | 0.2522 |
|  |  | Congruency | 0.02 | (2, 6237) | 0.9844 | 0.9882 |
|  |  | Age Group × Congruency | 0.84 | (2, 6238) | 0.4335 | 0.6503 |
|  | L Pos | Age Group | 0.14 | (1, 6071) | 0.7120 | 0.7932 |
|  |  | Congruency | 0.03 | (2, 6248) | 0.9672 | 0.9672 |
|  |  | Age Group × Congruency | 0.62 | (2, 6249) | 0.5409 | 0.7970 |
| R ATL | R Ant | Age Group | 4.33 | (1, 6122) | 0.0374 | 0.1123 |
|  |  | Congruency | 0.22 | (2, 6243) | 0.8004 | 0.8251 |
|  |  | Age Group × Congruency | 0.06 | (2, 6254) | 0.9439 | 0.9439 |
|  | R Pos | Age Group | 0.53 | (1, 6062) | 0.4671 | 0.7795 |
|  |  | Congruency | 0.05 | (2, 6252) | 0.9520 | 0.9520 |
|  |  | Age Group × Congruency | 1.34 | (2, 6263) | 0.2611 | 0.3916 |

**Note**. Abbreviations: *vmPFC*, ventromedial prefrontal cortex; *AG*, angular gyrus; *ATL*, anterior temporal lobe; *HPC*, hippocampus; *L*, left; *R*, right; *Ant*, anterior; *Pos*, posterior.
